# Myelin biogenesis is associated with pathological ultrastructure that is resolved by microglia during development

**DOI:** 10.1101/2021.02.02.429485

**Authors:** Minou Djannatian, Ulrich Weikert, Shima Safaiyan, Christoph Wrede, Cassandra Deichsel, Georg Kislinger, Torben Ruhwedel, Douglas S. Campbell, Tjakko van Ham, Bettina Schmid, Jan Hegermann, Wiebke Möbius, Martina Schifferer, Mikael Simons

## Abstract

To enable rapid propagation of action potentials, axons are ensheathed by myelin, a multilayered insulating membrane formed by oligodendrocytes. Most of the myelin is generated early in development, in a process thought to be error-free, resulting in the generation of long-lasting stable membrane structures. Here, we explored structural and dynamic changes in CNS myelin during development by combining ultrastructural analysis of mouse optic nerves by serial block face scanning electron microscopy and confocal time-lapse imaging in the zebrafish spinal cord. We found that myelin undergoes extensive ultrastructural changes during early postnatal development. Myelin degeneration profiles were engulfed and phagocytosed by microglia in a phosphatidylserine-dependent manner. In contrast, retractions of entire myelin sheaths occurred independently of microglia and involved uptake of myelin by the oligodendrocyte itself. Our findings show that the generation of myelin early in development is an inaccurate process associated with aberrant ultrastructural features that requires substantial refinement.

## INTRODUCTION

CNS myelination, a critical step in nervous system development, has enabled vertebrates to expand higher brain functions in a cost-efficient way (Stadelmann, Timmler, Barrantes-Freer, & Simons, 2019; Zalc, 2016). Myelin, produced by oligodendrocytes in the CNS, electrically insulates axons to facilitate saltatory conduction (Cohen et al., 2020; Hartline & Colman, 2007), while also providing metabolic support (Fünfschilling et al., 2012; Lee et al., 2012; Saab et al., 2016). Most of myelin is generated during early postnatal development according to a pre-defined genetic program. Once formed, myelin is a highly stable multilayered membrane, reflected by the unusually long lifetime of myelin-specific proteins (Toyama et al., 2013) and the low turnover of the sheaths. However, myelin remodeling can occur during learning and neuronal circuit refinement, thereby contributing to brain plasticity (Etxeberria et al., 2016; Fields, 2008; Gibson et al., 2014; Hill, Li, & Grutzendler, 2018; E. G. Hughes, Orthmann-Murphy, Langseth, & Bergles, 2018; Lakhani et al., 2016; Makinodan et al., 2012; McKenzie et al., 2014; Monje, 2018; Scholz, Klein, Behrens, & Johansen-Berg, 2009; Steele, Bailey, Zatorre, & Penhune, 2013; Wake et al., 2015; Xiao et al., 2016). In addition, a recent study in zebrafish demonstrated that developmental myelination is generated in excess and modified by microglial pruning (A. N. Hughes & Appel, 2020). We have previously provided a model of myelin wrapping, in which the leading edge moves around the axon underneath the previously deposited membrane, coupled with the lateral extension of myelin membrane layers along the axon (Snaidero et al., 2014). However, little is known about the error-rate and potential correction mechanisms that ensure proper myelin formation. Here, we investigated the ultrastructural changes of CNS myelin during development using three-dimensional serial block face scanning electron microscopy (SBF-SEM), focusing on abnormal myelination profiles. We show that myelin biogenesis is accompanied by the formation of abundant pathological ultrastructure. Phosphatidylserine, a known ‘eat-me’ signal, was identified on abnormal myelin profiles and on shedded myelin fragments. Live imaging in zebrafish showed that microglia phagocytose myelin and that myelin remodeling can also occur independently of microglia, in a process in which oligodendrocytes retract entire myelin sheaths into their cell body, followed by trafficking to lysosomes. Thus, our findings show that myelin biogenesis is error-prone requiring substantial refinement by oligodendrocytes and microglia.

## RESULTS AND DISCUSSION

### Myelin degeneration profiles occur during development and are resolved by microglia

To analyze myelin 3D ultrastructure in the developing CNS, we acquired volume stacks of 80 x 80 x 1150-210 µm size of P10, P14, P21 and P60 mouse optic nerves by SBF-SEM (Figure 1). As expected, myelination was sparse at P10 and sharply increased during the following two weeks, while all axons were myelinated by P60 (Figure 1A,C). Conversely, we found aberrant myelin structures in 24% of the myelinated axons on SEM cross sections at P10, which declined to 1.2% by P60 (Figure 1A,D; Video 1). These abnormal myelin profiles consist of redundant myelin outfoldings, whorls of degenerated myelin, split myelin, fragments that pinch off from a myelin sheath or are found in the extracellular space, and a so-far not described pattern that we term “myelin bulging” (Figure 1B,E; Video 1). Myelin bulging is characterized by an enlargement of the inner tongue and eccentric dilations of the myelin membrane, which partially folds back on itself, generating a myelin sheath that enwraps non-axonal structures (e.g., small processes of glial cells or even parts of another bulging myelin sheath; Figure 1F,G; Video 2). In a 3D volume, it is clearly distinguishable from double myelin, in which two myelin sheaths grow into each other (Djannatian et al., 2019). Bulgings were typically found adjacent to normal appearing myelin on the same sheath. While all types of myelin pathology decreased between P10 and P60, the fraction of myelin fragments was disproportionally large at P21 (49.1%). At P60, degenerated myelin and outfoldings were the most frequent pathologies (degenerated: 38.5%, outfoldings: 46.6%). Apart from the large whorl-like structures of degenerated myelin, we also observed focal dystrophies on normal-appearing myelin sheaths (Figure 2A, B). Both degenerated myelin and focal dystrophies were found to be engulfed by microglia, which often wrapped around several of these profiles at the same time (Figure 2A-C; Figure 2 − figure supplement 1; Video 3). Microglia also contained electron-dense fragments that likely represent products of myelin phagocytosis (Figure 2A, section 7). To confirm this, we co-immunostained tissue sections from mouse P14 and P80 corpus callosum (Figure 2C) and optic nerve (Figure 2D) with myelin-basic protein (MBP), IBA1 and LAMP1 to label microglia and late endosomes/lysosomes. About 3.5% of the microglia contained MBP^+^ fragments within LAMP1^+^ organelles at P10, while around 1.2% or less did at P80 (corpus callosum: P14: median 3.56% (interquartile range (IQR) 3.10-3.94%), P80: median 1.22% (IQR 0.93-2.27%), p=0.0286; optic nerve: P14: median 3.48%, (IQR 2.98-5.44%), P80: median 0.49% (IQR 0.31-1.32%), p=0.0286, Mann Whitney test, n = 4 mice). Together, these results show that microglia engulf and degrade myelin in mice under normal conditions. To clarify whether this role is unique to microglia, we performed immunostainings for astrocytes, which also have the capacity for phagocytosis previously described in the context of synapse elimination (Chung et al., 2013) and during metamorphosis(Mills et al., 2015). Indeed, we found MBP^+^ fragments in astrocytes of the corpus callosum (P14: median 2.08%, IQR 1.69-3.41%, P80: median 0.87%, IQR 0.50-1.38%, p=0.0286, Figure 2 – figure supplement 2).

**Figure 1.**
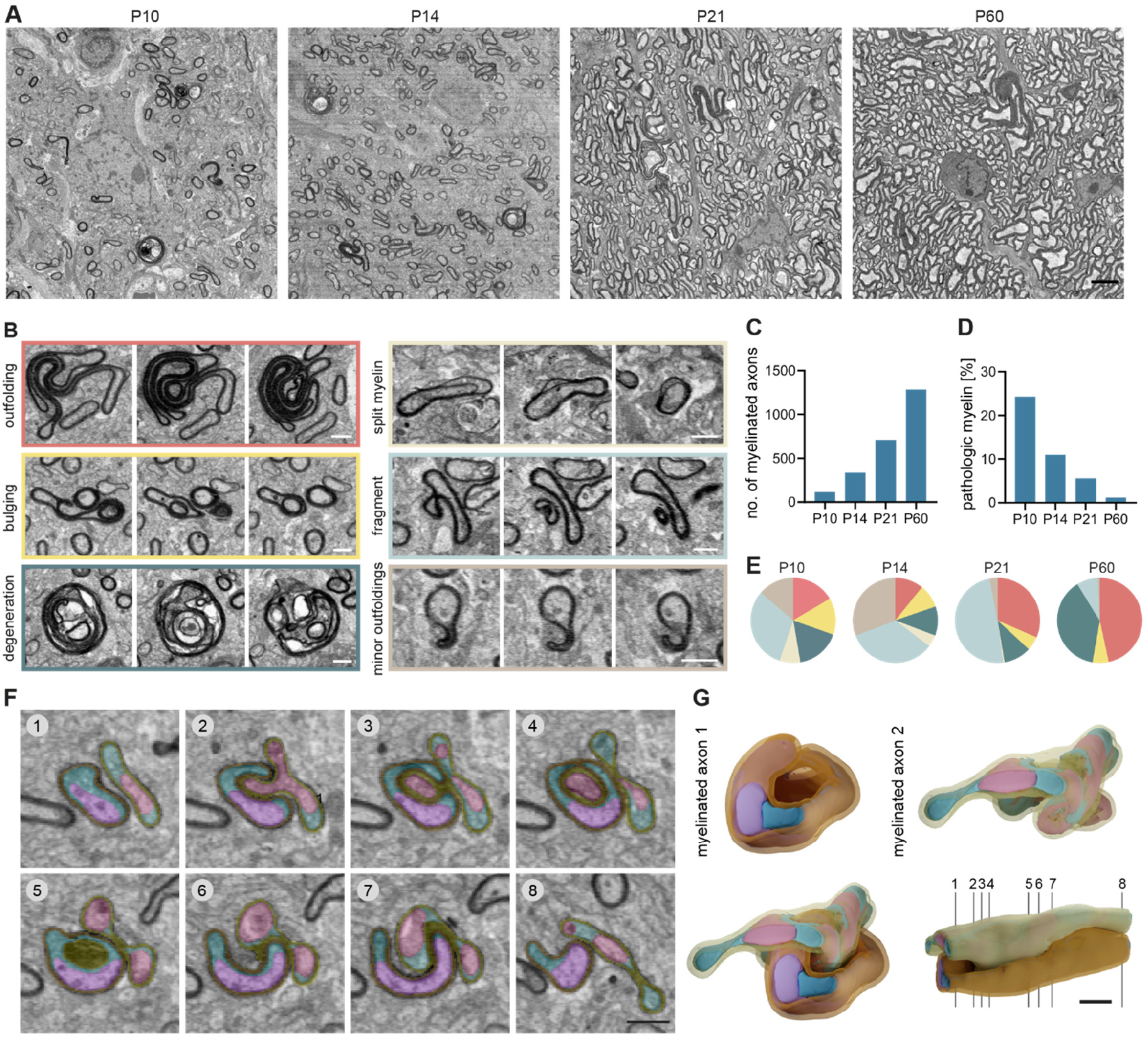
Myelin abnormalities occur temporarily during development in the mouse CNS. Serial block face scanning electron microscopy (SBF-SEM) of a P10, P14, P21 and P60 wild-type mouse optic nerves (80 x 80 x 150-210 µm volumes with a 10 x 10 x 80 nm resolution). (**A**) Representative cross-sections. (**B**) Cross-sections at different z levels show examples of myelin abnormalities: outfolding, bulging, degeneration, split myelin, fragment detaching from an intact sheath and minor outfoldings that likely represent an early stage of outfolding. (**C**) Quantification of the mean number of myelinated axons in a 40 x 40 µm area on 10 evenly dispersed cross-sections of the SBF-SEM volumes (n = 1 optic nerve). (**D**) Quantification of the mean number of myelin abnormalities within 10 evenly dispersed volumes (40 x 40 x 800 µm) normalized by the number of myelinated axons in reference sections (n = 1 optic nerve). (**E**) Distribution of myelin outfoldings (red), bulgings (yellow), degenerations (dark blue), split myelin (beige), fragments pinching off from a sheath or lying in the vicinity of a sheath (light blue) and minor outfoldings (brown) among abnormal myelin sheaths quantified in (D). (**F-G**) 3D reconstruction of a myelinated axon bulging into another myelinated axon. (F) Pseudo-colored cross-sections at different z levels (violet/pink: axons, petrol/cyan: glial cytoplasm, orange/yellow: myelin). Note the excess glial cytoplasm at the location of bulging. (G) 3D reconstruction. Numbering refers to cross-sections. Images in (A) and (G) are 4×4 binned. Scale bars: 3 µm (A), 2 µm (G), 1 µm (B, F). See Video 1 and 2.

**Figure 2 with 2 supplements.**
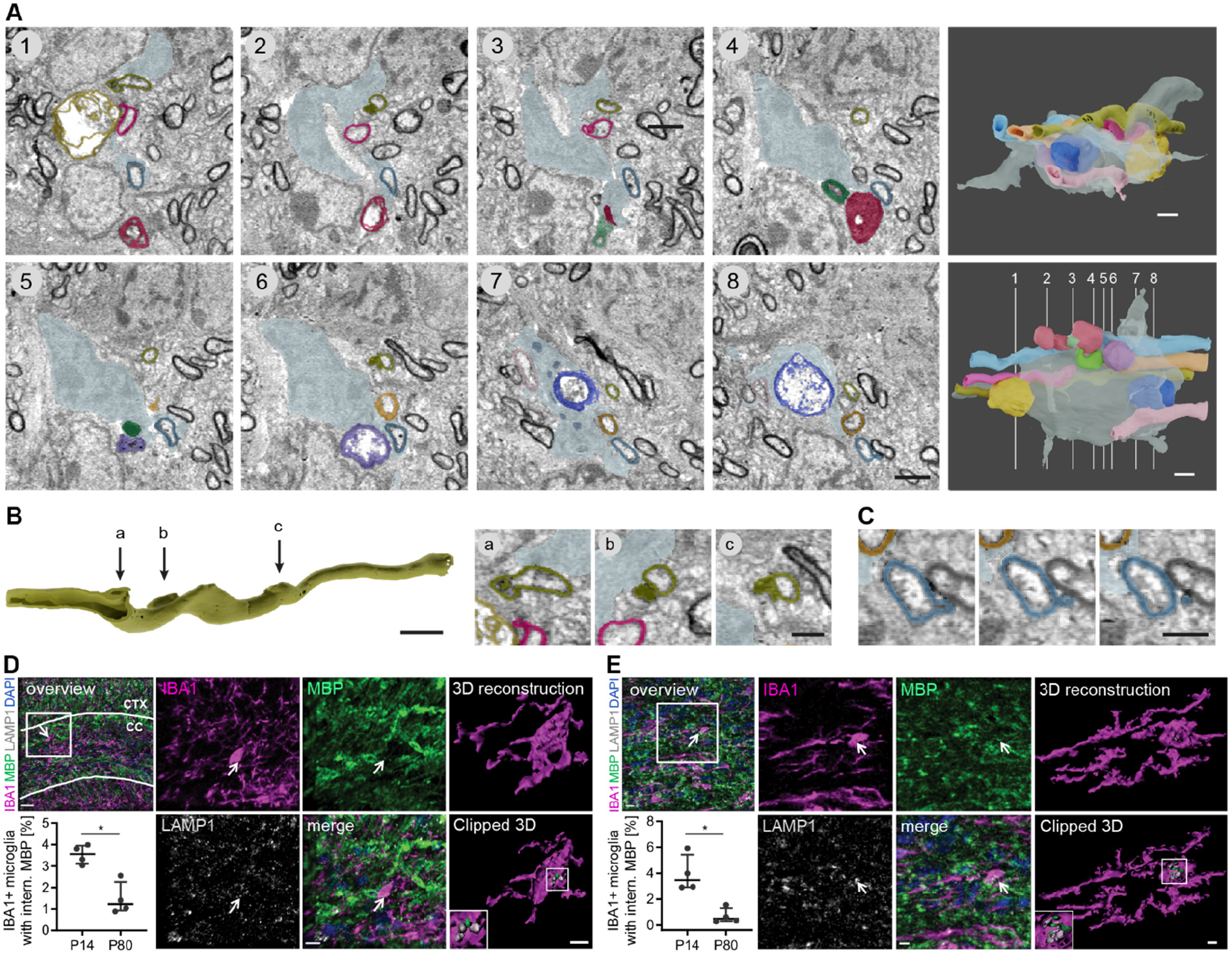
Microglia engulf and phagocytose developmental myelin pathology in the mouse CNS. (**A-C**) Serial block face scanning electron microscopy (SBF-SEM) of a P14 wild-type mouse optic nerve shows a microglia contacting several myelinated axons with pathologic myelin. (A) Pseudocolored cross-sections show examples of myelin pathologies engulfed or contacted by a microglia. The microglia is displayed in gray blue, while the other colors designate individual myelin sheaths. 3D reconstructions (right) show different orientations and numbering refers to cross-sections. (B) Focal myelin dystrophies on a reconstructed myelin sheath. The dystrophy in ‘a’ is sliced to display the 3D structure. Arrows refer to cross-sections on the right side. Cross-sections show details of the cross-sections 1,2 and 6 in (A). (C) Myelin fragment pinching off a myelin sheath on three consecutive sections. (**D-E**) Sections of P14 wild-type corpus callosum (D) and optic nerve (E) co-immunostained for MBP and LAMP1 together with IBA1 for microglia. Clipped 3D view shows mbp-positive staining inside microglia and in close association with lysosomes (arrows). Quantifications show the percentage of microglia with internalized MBP in P14 vs. P80 brains (n = 4 mice). Mann-Whitney U test: p= 0.0286 (C and D). Data represent median and interquartile range. Images in (A-C) are 4×4 binned. Scale bars: 30 µm (D, overview), 10 µm (D, ‘merge’, E, overview), 5 µm (D, clipped 3D, E, ‘merge’), 2 µm (A, B, 3D reconstructionsE, clipped 3 D), 1 µm (B, cross-sections, C). See also Figure 2 – figure supplement 1 and 2; Video 3.

### Microglia surveil and phagocytose myelin in zebrafish

In order to gain insight into how microglia interact with myelin *in vivo*, we conducted further experiments in the zebrafish spinal cord. This enabled us to follow myelination in a relatively short developmental window by expressing fluorescent reporters and implementing time-lapse imaging. When we imaged transgenic zebrafish larvae expressing mpeg1:EGFP (labelling microglia) and sox10:mRFP (labelling myelin) sheaths starting from the onset of myelination at 3 days post fertilization (dpf), we found that microglia were often elongated and in contact with myelin sheaths, (Figure 3A). During time-lapse imaging in 3-minute intervals over 1 hour, microglia extended and retracted their processes along myelin sheaths and, in parallel, moved their cell body forward (Figure 3 – figure supplement 1A; Video 4). In order to follow microglia movement along the spinal cord, individual microglia were followed in 20-30 minute intervals over 10-14 hours at 3-4 dpf (Video 4). Movements of individual microglia were tracked by their soma displacement in subsequent time frames (Figure 3B). Microglia moved with a mean velocity of 1.30 ± 0.59 µm/min (Figure 3C) and occasionally took breaks with negligible soma displacement over one or several subsequent time frames (average break duration: 72.00 ± 47.44 min, Figure 3D). We were able to identify distinct migrating patterns (Figure 3E): some microglia migrated from anterior to posterior (or vice versa) in a ‘linear’ fashion, while others performed a ‘circular’ movement, returning to approximately their original position within hours. We also observed combinations of these two patterns (‘undulating’) and microglia that remained at the same position during most of the acquisition (‘static’). The observation that some microglia returned to their previous position or remained static over longer periods prompted the question whether myelin sheaths in some oligodendrocyte territories are more intensely screened than others. We therefore analyzed microglial presence in quadrants of the dorsal and ventral spinal cord. Strikingly, we saw that microglial presence strongly deviated from a random distribution, with some quadrants being highly covered and others not at all during the entire acquisition (Figure 3F; Figure 3 – figure supplement 1C;; Video 4). Furthermore, high amounts of static phases were not compensated by a higher number of microglia (Figure 3 – figure supplement 1B).

**Figure 3 with 1 supplement.**
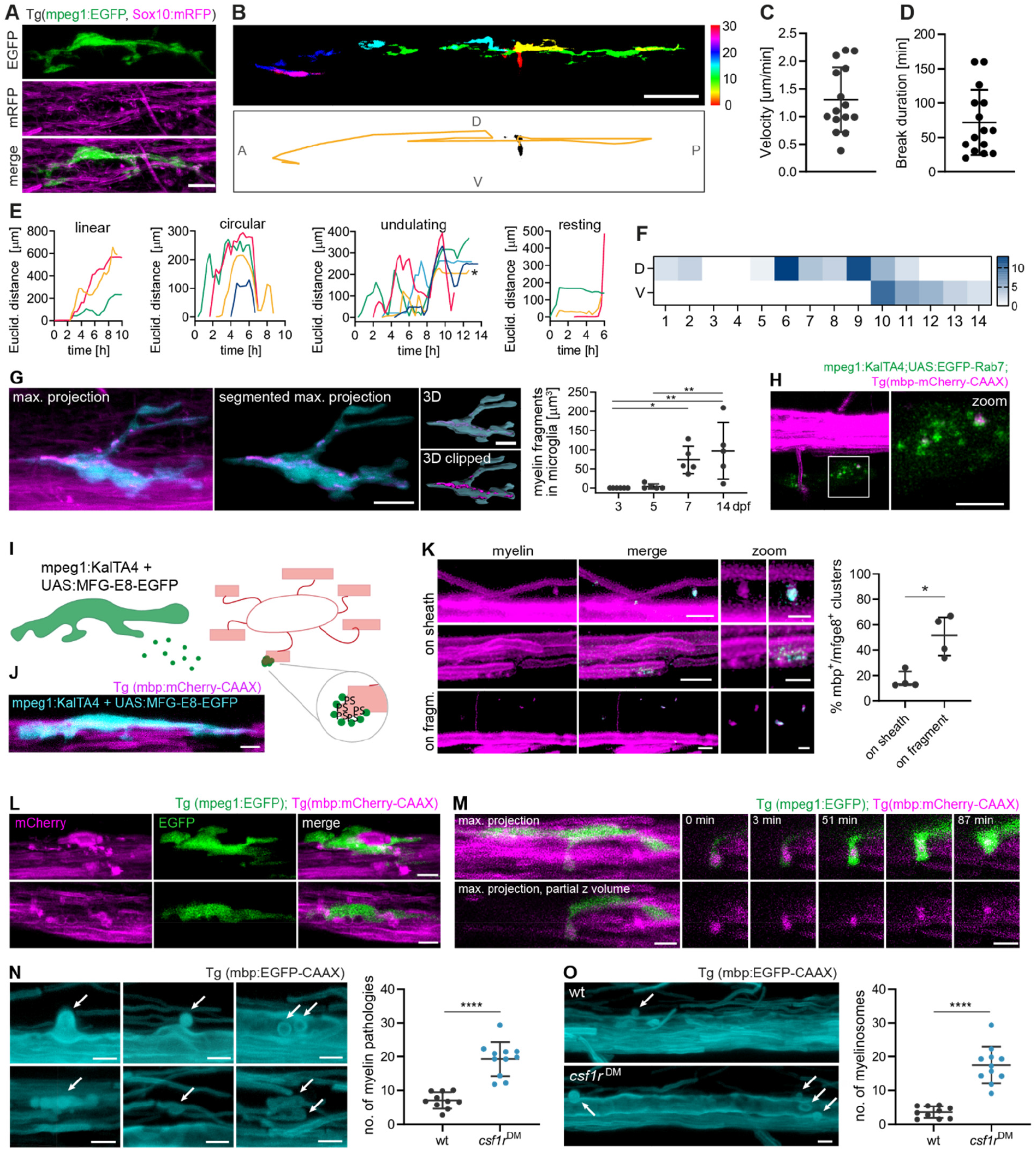
Microglia screen myelin sheaths and phagocytose myelin in the zebrafish spinal cord. (**A**) Maximum projection of a microglia (green) making contact with myelin sheaths (magenta) in a 4 dpf Tg(mpeg1:EGFP; sox10:mRFP) zebrafish spinal cord. (**B-F**) Microglia scanning behavior along myelin sheaths in the spinal cord was analyzed by confocal time-lapse imaging of 3-4 dpf Tg(mpeg1:EGFP; mbp:mCherry-CAAX) or Tg(mpeg1:EGFP; sox10:mRFP) larvae over 14.5-15 hours. Tile scans were acquired every 20 min. Individual microglia were followed through all time frames in which they were contacting myelin sheaths of one hemi-spinal cord (15 microglia from n = 6 larvae). (B) Example of the movement of a single microglia followed over 10 hours. *Top:* Individual time frames are color-coded to highlight microglia movement over time. *Bottom:* Binary image of the microglia in time frame 1. For the analysis of the microglia motility along the spinal cord, tracks were manually drawn in the Manual Tracking plugin in Fiji by marking soma position in each time frame (overlaid as a yellow line in this image). A: anterior, P: posterior, D: dorsal, V: ventral. (C) Mean velocity of microglia movement along the spinal cord). (D) Mean duration of breaks in motility taken by individual microglia. (E) Tracking profiles (as shown in B) were plotted as the Euclidean distance of the microglia soma in each frame from the position of the soma in the first frame. The shape of the resulting curves was used to classify microglia movements into patterns. Colors identify individual microglia. Asterisk marks sheath displayed in (B). (F) Microglia presence per time frame was added up in quadrants of the spinal cord (1 to 14, anterior to posterior) and displayed as heatmap to visualize preferential screening behavior. D: dorsal, V: ventral. (**G**) Myelin fragments inside of a microglia in a 10 dpf Tg(mpeg1:EGFP; sox10:mRFP) spinal cord. To quantify myelin fragments within microglia, microglia were manually segmented in Imaris and the resulting surfaces were used to mask the 552nm-channel. Quantification shows the sum of myelin fragment volumes within microglia in Tg(mpeg1:EGFP; mbp:mCherry-CAAX) spinal cords (n = 5 larvae per timepoint). One-way ANOVA with post-hoc Tukey’s test: 3 vs. 7 dpf: p=0.0345, 3 vs. 14 dpf: p=0.0046, 5 vs. 14 dpf: p=0.0089, all other comparisons were non-significant. (**H**) Colocalization of mbp:mCherry-CAAX^+^ fragments with microglial lysosomes, labelled by transiently expressed mpeg1:KalTA4;UAS:EGFP-Rab7. (**I**) Scheme depicting the *in vivo* detection of phosphatidylserine (PS), as performed in (J-K). Microglia secrete MFG-E8-EGFP, which binds to PS exposed on the extracellular leaflet of membrane. In compacted myelin membranes, extracellular exposure of PS is sterically hindered, but may occur upon pathological changes of the myelin sheath. (**J**) A microglial cell expressing MFG-E8-EGFP contacts mbp:mCherry-CAAX positive myelin sheaths. (**K**) Images show binding of MFG-E8-EGFP (cyan) to myelin sheaths or fragments (magenta) in Tg(mbp:mCherry-CAAX) larvae. Images are masked by the 552nm-channel. Zoomed images show details of myelin (left) and merged (right) images. Quantifications show the percentage of mbp:mCherry-CAAX^+^/MFG-E8-EGFP^+^ clusters on sheaths or on fragments. Mann-Whitney U test: p=0.0286. (**L**) Microglia engulfing an enlarged, seemingly unwrapping myelin sheath (*top row*) and surrounding round mbp-mCherry-CAAX-positive structures budding off from the ventral Mauthner axon (*bottom row*). (**M**) Confocal time-lapse imaging of a microglia taking up a myelin fragment over 90 minutes. Tilescans were acquired every 3 min. Images show maximum projections. For better visualization, only a part of the hemi-spinal cord was projected. Details show the myelin fragment inside the microglial process at distinct time points (top row: merged images, bottom row: 552 nm-channel). (**N**) Examples of myelin abnormalities observed in wild-type and *csf1r*^DM^ Tg(mbp:EGFP-CAAX) larvae (*top row*: myelinosomes budding off from the Mauthner axon, *bottom row left:* sheath degeneration, *bottom row middle + left:* “myelin flaps”, which may represent locally unwrapped myelin). Quantification of the number of myelin abnormalities in 10 dpf ventral spinal cord of wild-type (black) and *csf1r*^DM^ (blue) larvae, normalized by the myelinated area (wt: n = 10 larvae, *csf1r*^DM^: n = 11 larvae). Student’s t test: p< 0.0001. (**O**) Maximum projections of wild-type and *csf1r*^DM^ Tg(mbp:EGFP-CAAX) ventral spinal cords showing myelinosomes budding off from the Mauthner axon (arrows). Quantification of the number of myelin abnormalities in 10 dpf dorsal and ventral spinal cord of wild-type and *csf1r*^DM^ larvae (wt: n = 10 larvae, *csf1r*^DM^: n = 11 larvae).). Student’s t test: p< 0.0001. Data represent means ± s.d (C, D, G, N, O) and median and interquartile range (K). Scale bars: 2 µm (K, zoom), 5 µm (H-O), 10 µm (A, G), 50 µm (C). See also Figure 3 – figure supplement 1; Video 4.

We next asked whether microglia phagocytose myelin during their surveillance. Because the sox10:mRFP reporter transiently labels sensory neurons in addition to oligodendrocytes, we analyzed myelin phagocytosis in Tg(mpeg1:EGFP; mbp:mCherry-CAAX) animals. With time-lapse imaging, we observed the intake of a dense mbp:mCherry-CAAX^+^ myelin piece by a microglial process towards the microglial soma (Figure 3M; Video 4). At 3 dpf, no microglia contained myelin fragments, whereas all of them did by 7 dpf (n = 5 fish). Microglia showed increasing volumes of internalized mbp:mCherry-CAAX^+^ fragments between 5 and 7, or 14 days (Figure 3G). Importantly, microglia did not change their morphology towards an activated phenotype during myelin phagocytosis. Myelin fragments were highly motile inside microglia (Figure 3 – figure supplement 1D; Video 4) and co-localized with microglial lysosomes, as shown by the transient expression of the microglia-specific reporter mpeg1:KalTA4; UAS-EGFP-Rab7 (Figure 3H). Thus, microglia in zebrafish phagocytose myelin under physiological conditions, and accumulate it in lysosomes over time. These findings are consistent with a recent study by Hughes et al., which showed that myelin phagocytosis by microglia is dependent on neuronal activity (A. N. Hughes & Appel, 2020).

Next, we asked which molecular cues on myelin sheaths are recognized by microglia and promote myelin phagocytosis. Myelin is rich in the phospholipid phosphatidylserine (PS), and when exposed on the outer leaflet, it may serve as an ‘eat-me’ signal for microglia, which contain multiple PS receptors (Prinz, Jung, & Priller, 2019). To determine whether PS is exposed on degenerating myelin sheaths, we used a recently described PS reporter that can be applied *in vivo*. We took advantage of secreted glycoprotein MFG-E8, which specifically recognizes PS and, when fused to EGFP, can be delivered as a recombinant protein for PS detection (Kranich et al., 2020). Because injections of recombinant protein into the spinal cord may induce damage, we modified this approach by expressing the genetically encoded secreted MFG-E8-EGFP in microglia (Figure 3I,J). When we expressed mpeg1:KalTA4;UAS:MFG-E8-EGFP in transgenic mbp:mCherry-CAAX zebrafish larvae and masked MFG-E8-EGFP fluorescence by the mbp:mCherry-CAAX channel, we found that MFG-E8-EGFP localized to myelin sheaths (16.34 ± 6.58%) and to small, extracellular fragments positive for the myelin reporter (in 51.06 ± 16.15%) (Figure 3K). MFG-E8-EGFP-positive spots on myelin sheaths were focal and their inspection in 3D revealed that they marked small protrusions of the myelin sheath. We propose that these distinct irregularities on otherwise normal-appearing sheaths may eventually shed off, resulting in the release of extracellular fragments. These fragments may resemble “shedosomes” that occur during dendritic pruning (Han, Jan, & Jan, 2011) or “axosomes” which form during axon branch removal (Bishop, Misgeld, Walsh, Gan, & Lichtman, 2004).

We occasionally found microglia engulfing abnormal myelin sheaths that either lost their uniform 3D structure, or showed signs of fragmentation or local outfoldings (Figure 3L). However, myelin abnormalities were remarkably stable which made their removal by microglia difficult to follow with time-lapse imaging of larva beyond 5 dpf. We therefore asked whether myelin pathologies increase in the absence of microglia. For this, we analyzed a double mutant of the csf1 receptor, a duplicated gene in zebrafish, which is essential for microglia proliferation and development. These mutants maintain only around 5% of microglia in the CNS compared to wildtype animals (Oosterhof et al., 2018). Our characterization of the myelin in *csf1r*^DM^; Tg(mbp:EGFP-CAAX) larvae revealed mild hypomyelination (Figure 3 – figure supplement 1E,F) and a reduced oligodendrocyte number (Figure 3 – figure supplement 1G) in the spinal cord at 10 dpf, whereas myelin sheath length was unaffected (Figure 3 – figure supplement 1H). Interestingly, c*sf1r*^DM^ double mutant fish showed significantly more myelin pathologies in the 10 dpf ventral spinal cord compared to wild-type animals (wt: 5.5 ± 1.72, *csf1r*^DM^: 11.27 ± 2.76, p<0.0001, n = 10 (wt) and n = 11 (*csf1r*^DM^) fish; Figure 3N). These pathologies included fully degenerating sheaths and myelin segments that partially lost their 3D structure, making them appear like flaps. However, the increased myelin pathologies in *csf1r*^DM^ mutants were primarily due to the presence of bulb-like myelin outfoldings (Figure 3O), resembling ‘myelinosomes’ previously described in acute demyelinating EAE lesions(Romanelli et al., 2016).

### Oligodendrocytes retract myelin sheaths independently of microglia and degrade myelin membrane fragments

We also wondered whether myelin ingestion by microglia could be attributed to myelin sheath retractions (Auer, Vagionitis, & Czopka, 2018; Czopka, ffrench-Constant, & Lyons, 2013; Hines, Ravanelli, Schwindt, Scott, & Appel, 2015; Liu, Du, & He, 2013; Orthmann-Murphy et al., 2020; Yang, Michel, Jokhi, Nedivi, & Arlotta, 2020), a common form of myelin plasticity. We therefore analyzed retractions in 3-4 dpf Tg(mpeg1:EGFP; mbp:mCherry-CAAX) dorsal spinal cords in time-lapses of 30 minute intervals over 10-14 hours. The fine microglia processes that extended and retracted along myelin sheaths did not seem to be directly involved in any of the myelin retractions (Figure 4A; Video 5). In fact, microglial presence and the number of retractions along the spinal cord were entirely disconnected (Figure 4B), which we confirmed by a correlation analysis in 25 distinct territories of 1-2 oligodendrocytes, summed up over the entire acquisition (R^2^=0.0256, p=0.4453, n = 5 fish, Figure 4C). Furthermore, retractions did not significantly differ between microglia-depleted *csf1r*^DM^ and wild-type larvae (wt: 42.25 ± 7.50, *csf1r*^DM^: 57.75 ± 15.90, p=0.1429, Figure 4D; Video 5). Inspecting retractions more closely revealed that myelin sheaths were entirely withdrawn into oligodendrocyte cell bodies and no visible fragments were split off to the exterior. However, bright fragments appeared and shuttled within the oligodendrocyte soma in the course of retractions (Figure 4E; Video 5). We stained Tg(mbp:mCherry-CAAX) larvae with Lysotracker and analyzed the colocalization of these bright membrane fragments with lysosomes. We found that major portions of the fragments, but not the rest of the oligodendrocyte membrane, colocalized with lysosomes (Figure 4F). Intriguingly, we also found Lysotracker staining of myelin sheath pathologies, but not of normal-appearing adjacent sheaths or total myelin in the same image (Figure 4G, H), raising the possibility that some of the pathology is resolved by the oligodendrocyte itself. We next assessed whether myelin membrane fragments occur inside of the oligodendrocyte soma beyond the zebrafish model. For this, we co-immunostained mouse tissue sections of the optic nerve and the brain for MBP, the oligodendrocyte marker OLIG2, and LAMP1. In the corpus callosum, we found lysosome-associated MBP^+^ fragments in 0.53% (0.44-1.00%, median and IQR) of the OLIG2^+^ oligodendrocytes at P14, but only 0.20% (0.09-0.31%, median and IQR) at P80 (p=0.0286, Fig. 4I). Similarly, 1.92% (1.08-2.29%, median and IQR) of the oligodendrocytes in the optic nerve contained MBP^+^ fragments at P14, but only 0.30% (0.00-0.40%, median and IQR) did at P80 (p=0.0571, Figure 4I).

**Figure 4.**
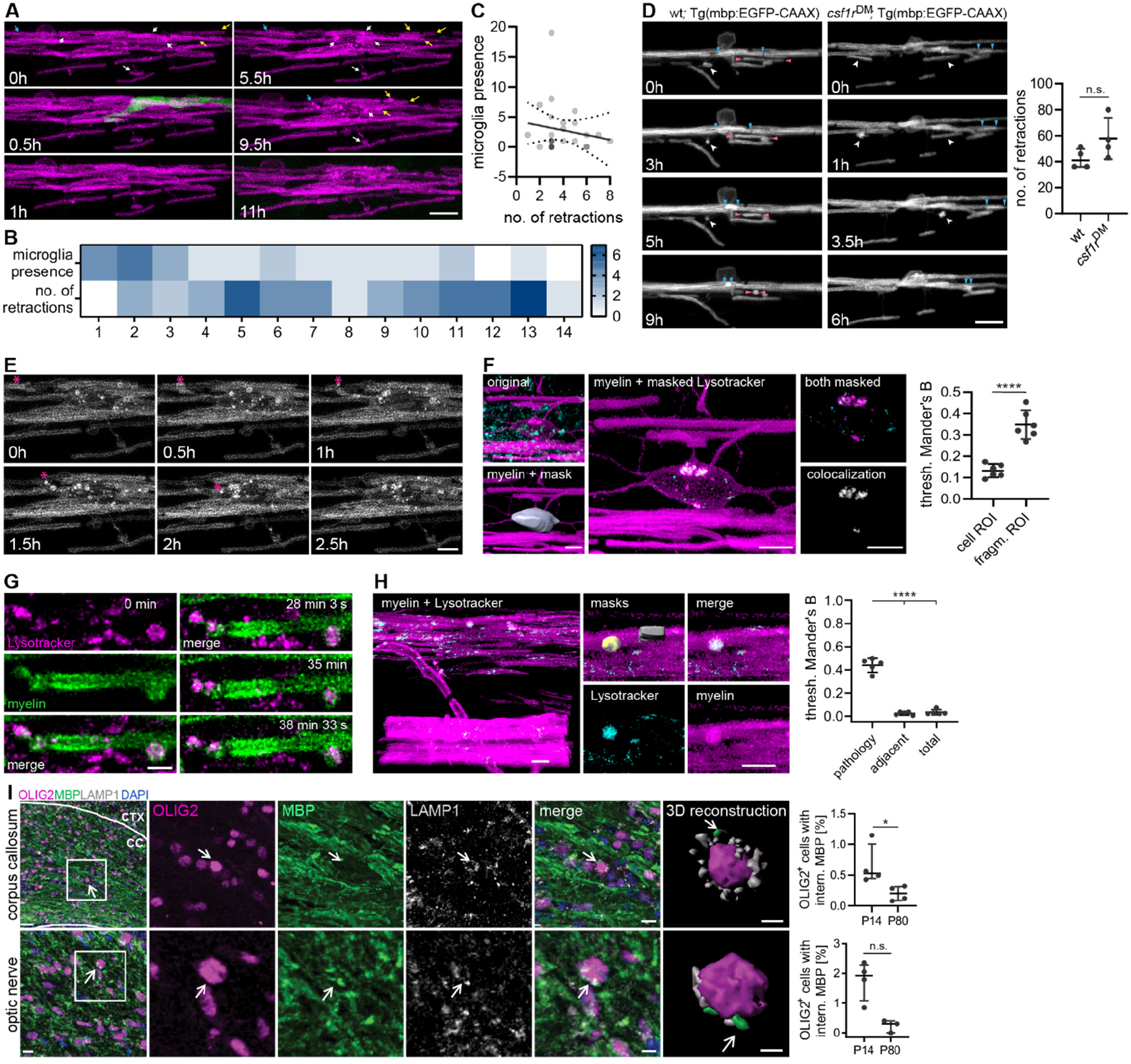
Oligodendrocytes retract myelin sheaths independent of microglia and degrade membrane fragments. (**A-C**) Time-lapse confocal imaging of 3 dpf wt Tg(mpeg1:EGFP; mbp:mCherry-CAAX) zebrafish spinal cord is used to analyze retractions of dorsal myelin sheaths with respect to microglial presence. Images were acquired every 30 min during 14 hours. Representative images are shown in (A). Arrows identify individual retractions by color and the position of the arrowheads. A microglia contacting myelin sheaths was only observed during a single frame (at 0.5 h). The heatmap in (B) shows microglial presence and the number of retractions in 5 x 5 mm quadrants along the dorsal spinal cord (1 to 12, anterior to posterior). XY plot in (C) correlates the number of time frames, in which a microglia was present, with the number of myelin sheath retractions in 26 oligodendrocyte territories (n = 5 larvae). Linear regression analysis: R^2^: 0.02556, p=0.4453. Dotted lines represent the standard error of the linear regression. (**D**) Time-lapse confocal imaging of 3 dpf wt and *csf1r*^DM^;Tg(mbp:EGFP-CAAX) larvae. Images were acquired every 30 min over 8 hours. Arrow heads identify individual retractions by different colors. Quantification shows the number of retraction during 8 hours. Mann-Whitney U test: p=0.1429. (**E**) Time-lapse confocal imaging from (A) shows how retractions result in bright fragments inside the oligodendrocyte cell body. Asterisk marks retracting sheath and its fragments. (**F**) Colocalization analysis of mbp:mCherry-CAAX-positive fragments inside oligodendrocyte cell bodies with Lysotracker Green. Oligodendrocyte cell bodies were manually segmented from the 552nm-channel. Channels were masked by the whole cell body (cell ROI) or by the bright mbp:mCherry-CAAX fragments inside (fragment ROI), followed by colocalization analysis. Thresholded Mander’s B coefficient reflects the amount of colocalization within the ROI of the thresholded 552nm-channel (43 cell bodies from n = 6 larvae). Student’s t-test: p< 0.0001. (**G**) Time-lapse imaging of a single myelin sheath stained in vivo with Lysotracker Red (maximum projections). Z-stacks were acquired every 3 s. (**H**) Quantification of lysotracker colocalization with myelin pathologies compared to a normal-appearing adjacent myelin sheath and total myelin in the same image (n = 5 larvae). One-way ANOVA with post-hoc Tukey’s test: pathology vs. adjacent, pathology vs. total: p<0.0001, adjacent vs. total: p=09074. (**I**) Free-floating sections of P14 wild-type mouse brains co-immunostained for MBP and LAMP1 together with OLIG2 for oligodendrocytes. 3D reconstruction shows mbp-positive staining in close association with lysosomes and the oligodendrocyte nucleus. Quantifications show the percentage of oligodendrocytes with internalized MBP in P14 vs. P80 brains (n = 4 mice). Student’s t test: p=0.0286 (corpus callosum), p=0.0571 (optic nerve). Data represent means ± s.d (E, H) and median and interquartile range (D, I). Scale bars: 30 µm (I, corpus callosum overview), 10 µm (A, D, I, optic nerve overview and corpus callosum ‘merge’), 5 µm (E, F, H, I, optic nerve ‘merge’), 3 µm (clipped 3Ds), 2 µm (G). See also Video 5.

In summary, we have shown that CNS myelin undergoes significant ultrastructural changes during development that are almost entirely resolved by adulthood. Ultrastructural analysis and time-lapse imaging revealed extrinsic and intrinsic mechanisms of myelin clearance. Extrinsic myelin removal includes the clearance of degenerated myelin and shedded myelin fragments by microglia and, possibly, astrocytes. Exposed PS on myelin presents a phagocytic signal for microglia, which express several PS-mediated receptors (Lemke, 2019). In an intrinsic mechanism related to neuronal activity (Hines, Ravanelli, Schwindt, & Scott, 2015; E. G. Hughes et al., 2018; Mensch et al., 2015), oligodendrocytes remove whole myelin sheaths by retractions into their cell body and lysosomal degradation therein. These processes can collectively result in a refinement of how myelin supports neuronal circuits while ensuring its proper function. Previous studies have shown that myelin remodeling occurs during learning and neuronal circuit refinement (Etxeberria et al., 2016; Gibson et al., 2014; Hill et al., 2018; E. G. Hughes et al., 2018; Lakhani et al., 2016; Makinodan et al., 2012; McKenzie et al., 2014; Monje, 2018; Scholz et al., 2009; Steele et al., 2013; Xiao et al., 2016). This is associated with modifications of myelin sheath length and thickness, the addition of new myelin segments along an axon and retractions of whole sheaths (Auer et al., 2018; Etxeberria et al., 2016; Ford et al., 2015; Kaller, Lazari, Blanco-Duque, Sampaio-Baptista, & Johansen-Berg, 2017; Sinclair et al., 2017). It is tempting to speculate that the failure to clear degenerated myelin and/or a dysbalance of myelin homeostasis during development leads to nervous system dysfunction and neurocognitive disorders. Intriguingly, the onset of psychiatric disease in young adulthood typically coincides with a phase of enhanced myelin plasticity (Höistad et al., 2009; Miller et al., 2012). Furthermore, developmental myelin abnormalities, changes in myelination patterns and microglial hyperactivity have been linked to the pathogenesis of schizophrenia (Janova et al., 2018; Kelly et al., 2018; Raabe et al., 2019; Sellgren et al., 2019; Takahashi, Sakurai, Davis, & Buxbaum, 2011; Uranova, Vikhreva, Rachmanova, & Orlovskaya, 2011). Future studies need to investigate the possible contribution of myelin remodeling and clearance in neuropsychiatric diseases.

## MATERIALS AND METHODS

### Key resources table

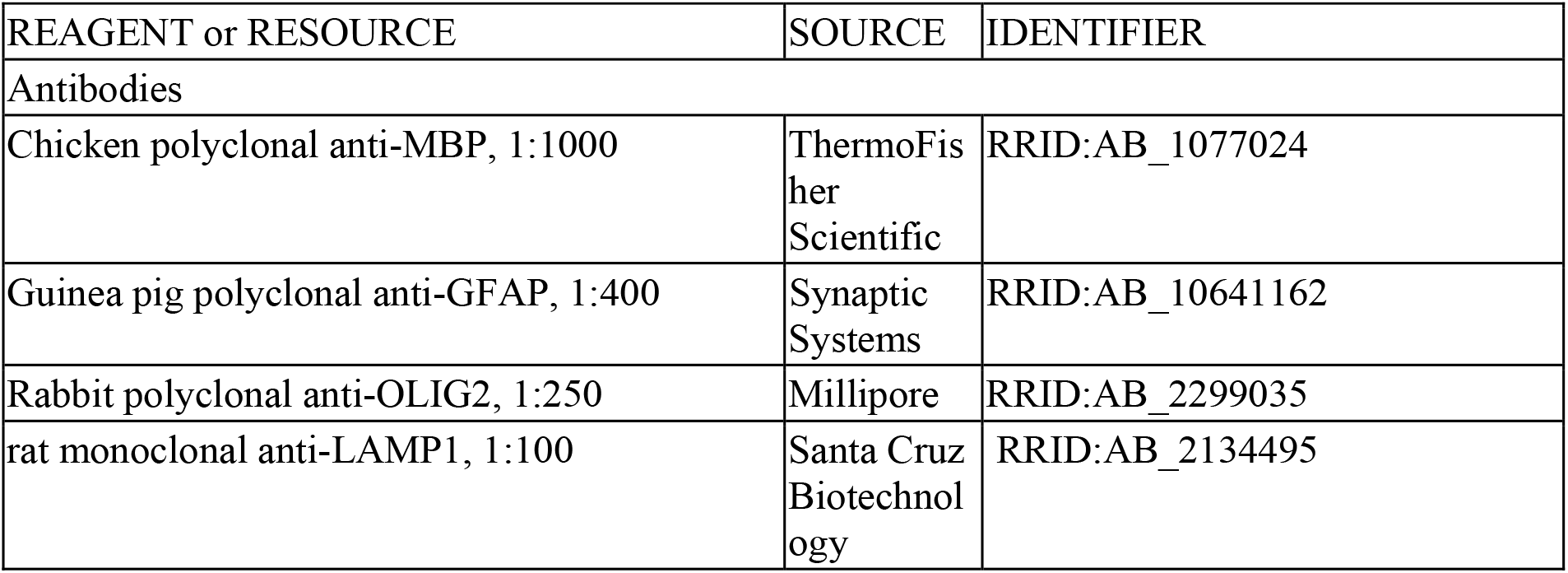
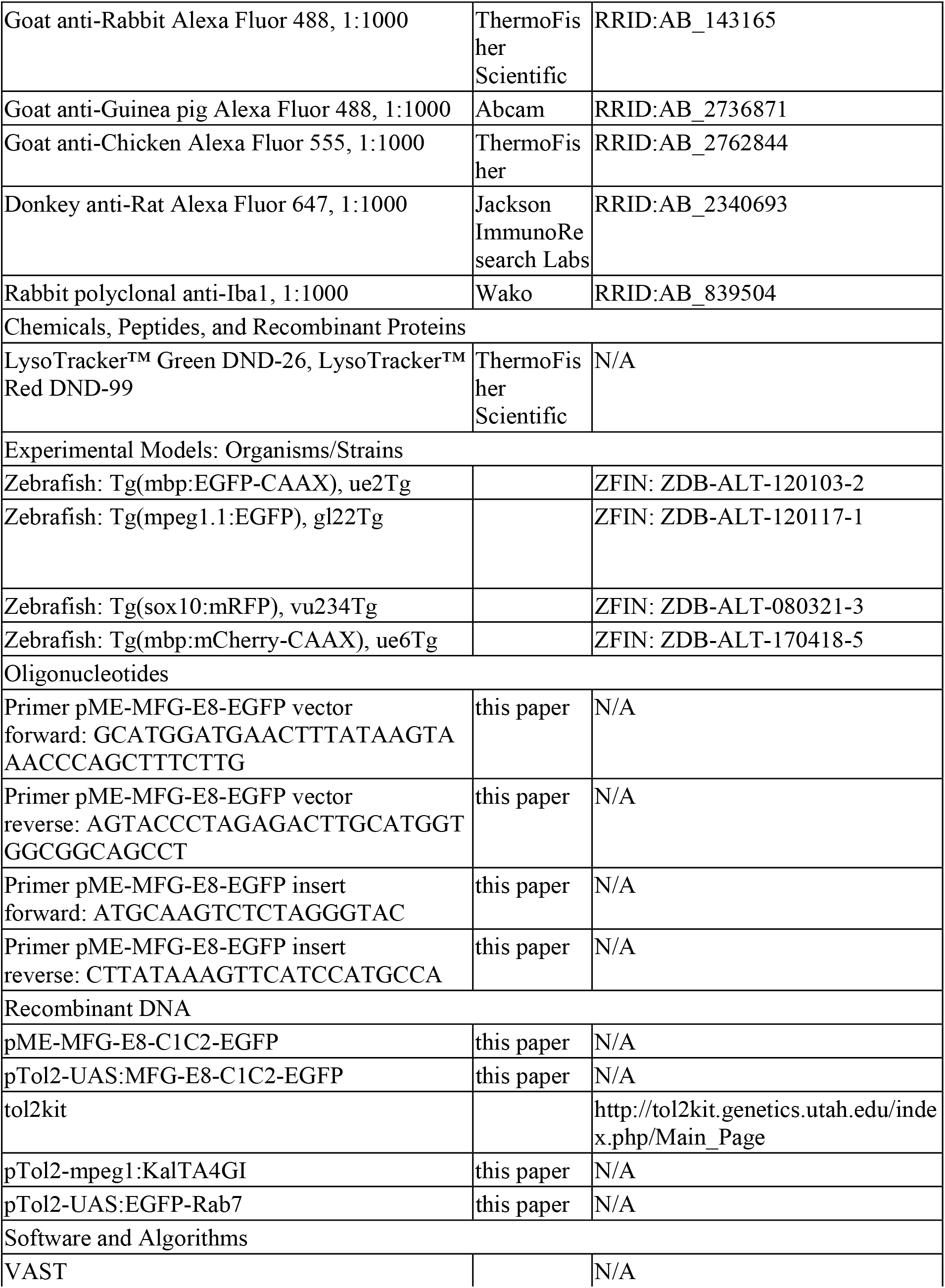
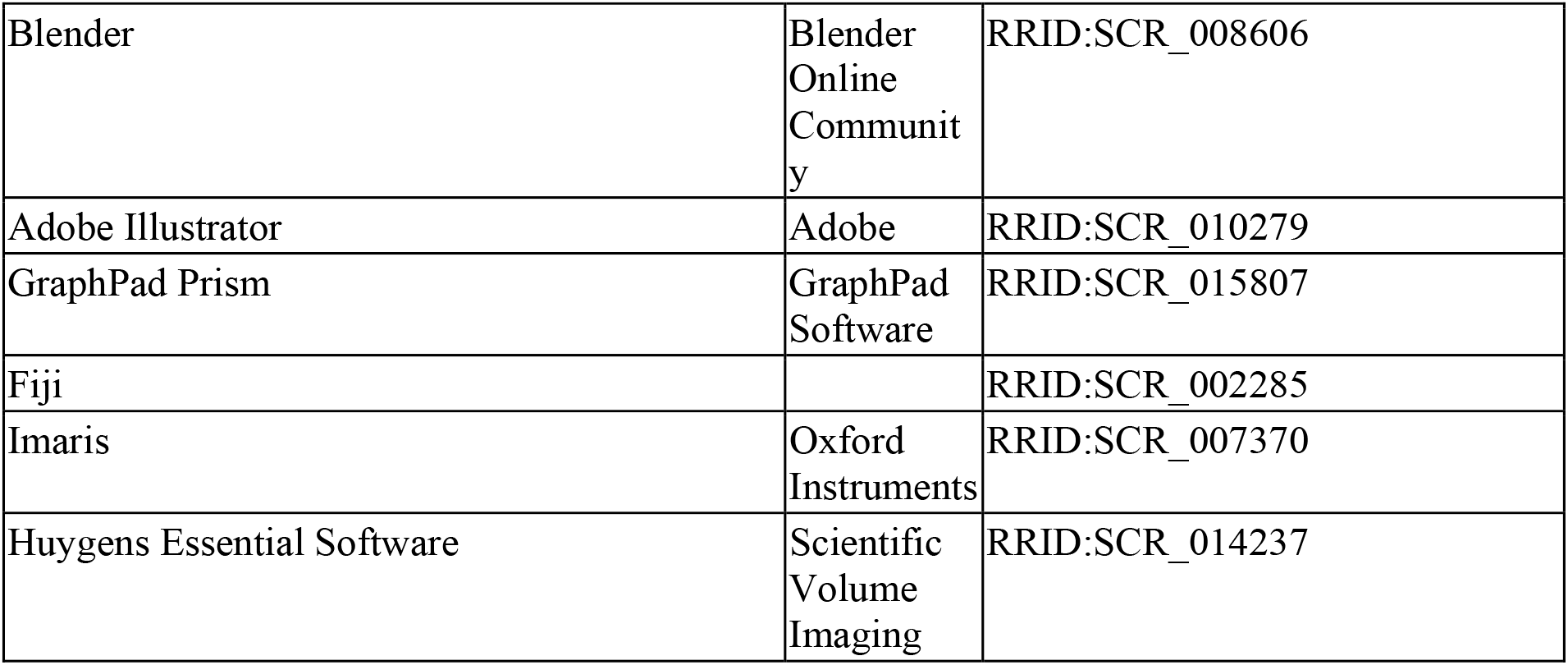

### Zebrafish husbandry

We used the existing transgenic zebrafish lines Tg(mpeg1:EGFP), Tg(sox10:mRFP) and Tg(mbp:mCherry-CAAX) as well as the mutant line *csf1r*^DM^ (containing mutant alleles *csf1ra* and *csf1rb*) which was previously outcrossed to Tg(mbp:EGFP-CAAX). All zebrafish procedures were carried out with approval and according to the regulations of the District Government of Upper Bavaria. Zebrafish were housed at the fish facility of the German Center for Neurodegenerative Diseases (DZNE) in Munich according to local animal welfare regulations. Embryos were obtained by natural spawning and raised at 28.5°C in E3 medium.

### Mouse husbandry

Male and female C57BL/6 mice were group housed in a 12-hour dark/light cycle with ad libitum access to food and water in the animal facility of the Max Planck Institute of Experimental Medicine in Göttingen. All animal procedures were handled with approval and in accordance to the regulations of the state government of Lower Saxony.

### Generation of transgenic constructs

To generate pTol2-mpeg1:KalTA4, we first cloned the middle entry vector pME-KalTA4GI by a BP reaction of pTol2-sox10:KalTA4GI (kind gift of Tim Czopka(Almeida & Lyons, 2015)) with pDONR 221, using Gateway™ BP Clonase™ II Enzyme mix (ThermoFisher Scientific). pME-KalTA4GI was then recombined with p5E-mpeg1 and p3E-polyA into pDestTol2CG2(Kawakami, 2007), using LR Clonase™ II Plus enzyme (ThermoFisher Scientific). pTol2-UAS:EGFP-Rab7 was generated by recombining p5E-UAS, pME-EGFP no stop and p3E-Rab7 (kind gift of Brian Link(Clark, Winter, Cohen, & Link, 2011)) into pDestTol2-CG2 in an LR reaction, as described above. To obtain pTol2-UAS:MFG-E88-C1C2-EGFP, we first cloned pME-MFG-E8-C1C2-EGFP by PCR amplification of the mouse MFG-E8 C1 and C2 domain (insert) and of the middle entry vector backbone (vector), using Q5 polymerase (New England Biolabs Inc). Fragments were generated with the following primers (template plasmids indicated in brackets): insert fragment (pD2523-mMFG-E8_C1C2-EGFP, kind gift of Jan Kranich(Kranich et al., 2020)): 5’-ATGCAAGTCTCTAGGGTAC-3’ (fwd) and 5’-CTTATAAAGTTCATCCATGCCA-3’ (rev), vector fragment (pME-KalTA4GI): 5’-GCATGGATGAACTTTATAAGTAAACCCAGCTTTCTTG-3’ (fwd) and 5’-AGTACCCTAGAGACTTGCATGGTGGCGGCAGCCT-3’ (rev). This was followed by a 2-fragment Gibson assembly, using the NEBuilder® HiFi DNA Assembly Cloning Kit (New England Biolabs, Inc.) according to the manufacturer’s protocol. pTol2-UAS:MFG-E8-C1C2-EGFP was then generated by recombining p5E-UAS, pME-MFG-E8-C1C2-EGFP and p3E-polyA into pDestTol2-CG2 in a Gateway LR reaction. All sequences were verified by Sanger sequencing.

### Microinjection of zebrafish embryos

Transgenic reporter constructs were transiently expressed in zebrafish by injecting a mixture of plasmid DNA (25 ng/μl each), transposase mRNA (25 ng/μl) and 0.2 M KCl into fertilized eggs at the one-cell stage. Embryos for imaging were treated with PTU from 8–24 hpf onwards to prevent pigmentation.

### Lysotracker staining in zebrafish

Zebrafish larvae were incubated for 2 hours with 1:100 LysoTracker™ Green DND-26 or LysoTracker™ Red DND-99 (ThermoFisher Scientific) in E3 medium (final concentration of 10 µM). Thereafter, larvae were washed three times with E3 medium, mounted and immediately imaged.

### Immunohistochemistry

30μm free-floating brain sections and 14μm mounted optic nerve sections were rinsed with 1x PBS containing 0.2% Tween-20 and permeabilized in 0.5% Triton X-100 for 30 min. Sections were blocked for 1 h at room temperature in a solution containing 2.5% FCS, 2.5% BSA and 2.5% fish gelatin in PBS. Primary antibodies, diluted in 10% blocking solution, were added and incubated overnight at 4°C. On the following day, sections were incubated with secondary antibodies, diluted in 10% blocking solution, for 1 h at room temperature. After washing with PBS, the free-floating sections were mounted on superfrost plus slides using fluorescence mounting medium (Dako). Primary antibodies: rabbit IBA1 (Wako, Cat., 019-19741, 1:1000), chicken MBP (ThermoFisher Scientific, Cat, PA1-10008, 1:1000), guinea pig GFAP (Synaptic Systems, Cat., 173004, 1:400), rabbit Olig2 (Millipore, Cat., AB9610, 1:250), rat LAMP1 (1D4B) (Santa Cruz, sc-19992, 1:100). Secondary antibodies: Alexa Fluor 488, 555 and 647-conjugated antibodies (ThermoFisher, Abcam, Jackson ImmunoResearch Labs, 1:1000).

### Confocal image acquisition

Zebrafish larvae at 3 to 14 dpf were anesthetized with tricaine and mounted laterally in 1% low melting point agarose (ThermoFisher) on a glass bottom dish (#1.5 cover glass, IBL). Fish were imaged at a Leica TCS SP8 confocal laser scanning microscope with automated moving stage and climate chamber (28.5°C), using a resonant scanner, 1.1 NA 40× water immersion objective and 488 and 552 nm lasers. Single images or tilescans (1248×1248 pixels, pixel size 40-78 nm, z step size 250-320 nm) were acquired using a hybrid detector in counting mode, line accumulation of 4-8, and the pinhole at 0.8 airy units. Overnight time lapse imaging (800-1248 × 800-1248 pixels, pixel size 78-120 nm, z step size 0.5-1 sµm) was done with a hybrid detector in photon counting mode, line accumulation of 4-8, and a pinhole at 1.0-1.2 airy units. z-stack tiles (z-step = 1 µm) along the spinal cord, starting at the neck, were taken every 20-30 min. Time-lapses were registered with the Correct 3D drift plugin in Fiji. Images were stitched using the LAS X (v3.5) software. For better representation and to improve colocalization analysis, images showing fine details or colocalizations (Figs. 3A, 3G-H, 3K-M, 4A, 4E-H and Suppl. Fig. 3D) were deconvolved using the default settings of the Huygens Essential software (Scientific Volume Imaging). All images of zebrafish show lateral views of the spinal cord with anterior to the left and dorsal to the top. Images of mouse immunostainings were acquired via a Leica TCS SP5 confocal microscope and were processed and analyzed with Imaris (64x version 9.2.0). To estimate the number of Iba1 positive cells with internalized myelin particles, confocal stacks (step size: 0.8μm) were captured in the z-direction from the region of interest (corpus callosum and optic nerve) with 20X objective of the confocal microscope. A three dimensional view was created from stacks of each image using the surpass option in Imaris.

### Tissue preparation for SBF-SEM and image acquisition

Animals were sacrificed by cervical dislocation and the optic nerves were dissociated from the eyeballs and the chiasm. They were immersion fixed for 24 h in Karlsson Schultz solution (4% PFA, 2.5% glutaraldehyde in 0.1M PBS (pH 7.3)) (Karlsson & Schultz, 1965) at 4°C. Afterwards, they were processed with a modified reduced osmium-thiocarbohydrazide-osmium (rOTO) protocol (Mikula, Binding, & Denk, 2012) to increase the contrast. At first they were washed 3 times with 0.1M PB buffer followed by 3 h of 2% OsO4 and 0.25% K4[Fe(CN)6] incubation at 4°C to reduce OsO4 to OsO2. After a wash step in ddH20, samples were incubated with 0.1% thiocarbohydrazide (in ddH2O) for 1 hour at room temperature. Contrast was achieved by incubation with 2% OsO4 for 90 minutes followed by (after washing with ddH2O) 2.5% uranyl acetate incubation over night at 4°C. After washing again in ddH2O, samples were dehydrated using increasing concentrations of acetone (always 15 minutes at 30%, 50%, 75%, 90%, 3x 100%). Samples were embedded using an increasing concentration of Durcupan resin (without component D) in acetone (1.5 hours at 25%, 50%, 75%, and overnight at 100% Durcupan), followed by incubation with 100% Durcupan + component D for 5 hours and final embedding in silicone molds with fresh resin from the same composition. They were polymerized for 48 hours at 60°C until the blocks hardened. The sample in the resin was exposed over a block length of ∼400 μm on four sides by trimming away the excess resin using a trimming diamond knife (Ultratrimm 90°, Diatome, Biel, Switzerland). The trimmed block was mounted on an aluminum Gatan pin with CW2400 conductive epoxy resin and baked for additional 3 days at 60°C. Finally, the block was sputter coated with 20 nm of platinum (ACE600 Leica Microsystems, Vienna) before imaging at the Zeiss Merlin VP Compact SEM equipped with a Gatan 3View2XP system. SEM images of the block-face were acquired at the variable pressure mode by an alternating procedure of cutting 80 nm and imaging with a pixel time of 0.5 μs and an image size of 8000 x 8000 pixels with 10 nm pixel size. About 1900-2700 sections per block were acquired.

### 3D reconstruction of SEM images

Volumes were 4×4 binned for 3D image reconstruction. Cropped volumes containing the regions of interest were aligned with the TrakEM plugin (Cardona et al., 2012) of Fiji. Aligned stacks were manually segmented in VAST (Berger, Seung, & Lichtman, 2018) and renderings were done using the Blender software.

### Quantifications

#### Analysis of mouse optic nerve SBF-SEM

The total number of myelinated axons was counted on ten 40 x 40 µm sections (reference sections). Structural defects were assessed within small volumes of in total 100 sections around the reference section and normalized by the total number of myelinated axons. Counting was done with the Cell Counter tool in Fiji.

#### Analysis of myelin fragments in mouse immunohistochemistry

The number of cells with internalized myelin particles, labeled by anti-MBP and colocalizing with anti-LAMP1, was quantified and normalized by the total numbers of cells in an area of 0.6 mm^2^.

#### Tracking of microglia motility in zebrafish

Tracking of microglia soma displacement was done with the Manual Tracking plugin in Fiji (F. Cordelières, Institut Curie, Orsay). Velocity between two consecutive time frames was an output of the Manual Tracking plugin and mean velocity represents the mean velocity of an individual microglia over all time frames. A time frame was defined as a break when no soma displacement had taken place compared to the previous frame. Break duration was calculated for individual microglia by adding consecutive time frames in which breaks occurred, multiplied by the acquisition interval. Euclidean distances were calculated from X,Y coordinates recorded by the plugin. For heatmaps, a 5 x 5 mm grid was placed over the spinal cord, separating the dorsal and ventral spinal cord. Presence of a microglia (or parts of it) in each quadrant were added up in all time frames.

#### Analysis of csf1r^DM^ mutants

Analysis was performed blinded for genotypes. Myelinated area in the ventral or dorsal spinal cord was determined in Fiji as the total area of mbp:EGFP-CAAX-positive pixels from ROIs of manually thresholded maximum intensity projections. Number of oligodendrocytes, sheath lengths, and myelin pathologies were quantified using the 3D surpass view of IMARIS (Bitplane). For sheath lengths, distances between sheath ends were measured in the dorsal spinal cord, including sheaths of commissural neurons that could be followed along their entire length. Myelin pathologies were assessed in the 3D view of Imaris. The number of myelin pathologies and myelinosomes was normalized by the myelinated area in the ventral spinal cord.

#### Quantification of myelin sheath retractions

Myelin sheaths in the dorsal spinal cord showing negative growth over consecutive time frames were quantified as retractions. These were assessed on maximum intensity projections and verified on z-stacks in Fiji. For the heatmap, the spinal cord was divided into 5 x 5 mm quadrants. Per quadrant, we quantified the sum of time frames in which a microglia was present and the number of retractions over the whole acquisition. We did a similar quantification to assess the correlation between microglial presence and retractions, but on the basis of oligodendrocyte territories (containing the myelin sheaths made by 1-2 oligodendrocytes) instead of quadrants.

#### Volume of myelin fragments inside microglia

Microglial volumes were manually segmented in Imaris and the resulting surfaces were used to mask both channels (488 nm-channel for microglia and 552 nm-channel for myelin). The binary masked 488 nm-channel was then despeckled in Fiji to remove random noise. Microglia and myelin volumes were calculated with the Voxel Counter plugin in Fiji. Data are represented as means per individual larvae.

#### Colocalization of LysoTracker staining with myelin fragments inside oligodendrocytes

Oligodendrocyte cell bodies were manually segmented in Imaris and the resulting surfaces were used to mask both channels as “cell ROI” (488 nm-channel for LysoTracker and 552 nm-channel for myelin fragments). A second surface encompassing the bright myelin fragments within oligodendrocyte cell bodies was created by thresholding the first surface by intensity. Masking of both channels by this surface resulted in “fragment ROIs”. Masked channels were manually thresholded for colocalization analysis in ImarisColoc. Thresholded Mander’s B coefficients represent the colocalized fraction within the myelin channel after thresholding. Data are represented as means per individual larvae.

#### Colocalization of LysoTracker staining with myelin pathology

Myelin pathologies and parts of adjacent myelin sheaths were manually segmented in Imaris. Additionally, an automatic surface was created for the 552 nm-channel, representing total myelin sheaths (and membranes of oligodendrocytes somas). The resulting surfaces from these three segmentations were used to mask both channels (488 nm-channel for LysoTracker and the 552 nm-channel). Masked channels were manually thresholded in ImarisColoc using the same thresholding parameters for myelin pathologies, adjacent sheaths and total myelin. Colocalization analysis for these three categories was carried out in ImarisColoc. Thresholded Mander’s B coefficients represent the colocalized fraction within myelin. Data are represented as means per individual larvae.

#### Analysis of mbp:mCherry-CAAX^+^/MFG-E8-EGFP^+^ clusters

mbp:mCherry-CAAX^+^/MFG-E8-EGFP^+^ clusters were assessed on images masked by the 552 nm-channel by a surface automatically created in Imaris. Clusters were counted in the 3D view in Imaris. Data are represented as means per individual larvae.

#### Statistical analysis

Statistical analysis was performed in GraphPad Prism. All samples were tested for normality. Comparison of two independent groups was carried out using student’s t test (for normally distributed data) and Mann-Whitney U test (for non-normally distributed data). Groups with small sample sizes that were visibly non-normally distributed were tested by Mann-Whitney U test. Comparisons between three or more groups were done by one-way ANOVA, followed by Tukey’s post-hoc test for multiple comparisons. Data in the text is presented as mean ± s.d. or median and interquartile range (25^th^ percentile - 75^th^ percentile) as indicated in the legends. Correlations were tested by linear regression and the coefficient of determination (R^2^) and p value are reported in the legends. P values are indicated throughout the figures as follows: n.s.: p≥ 0.05, *: p< 0.05, **: p< 0.01, ***: p< 0.001, ****: p< 0.0001.

## ACKNOWLEDGMENTS

We thank Tim Czopka, David Lyons, Brian Link, Jan Kranich, Francesca Peri and Michel Reimer for providing reagents.

## ADDITIONAL INFORMATION

### Author contributions

M.S. and M.D. designed the project. M.D., U.W, S.S., C.W., C.D., G.K., T.R. and D.C. performed experiments and/or analyzed the data. T.v.H., M.Sch., W.M., J.H., B.S. and M.S. analyzed the data or supervised data acquisition. M.D. wrote the first draft. M.S. and M.D. edited the manuscript with input from all authors.

### Ethics statement

All zebrafish procedures were carried out with approval and according to the regulations of the District Government of Upper Bavaria, Germany. All mouse procedures were handled with approval and in accordance to the regulations of the state government of Lower Saxony, Germany.

## ADDITIONAL FILES

### Supplementary files

**Video 1:** Myelin pathologies are abundant during development and are resolved until young adulthood.

**Video 2:** 3D reconstruction of a myelin bulging.

**Video 3:** Microglia are associated with myelin pathologies.

**Video 4:** Myelin screen myelin along the spinal cord and phagocytose myelin debris.

**Video 5:** Myelin retractions occur independently from microglia and are broken down into fragments in the oligodendrocyte cell body.

### Data availability

All measurements and statistical analyses generated and used in this study are included in the manuscript.

---

**Figure 2 – figure supplement 1.**
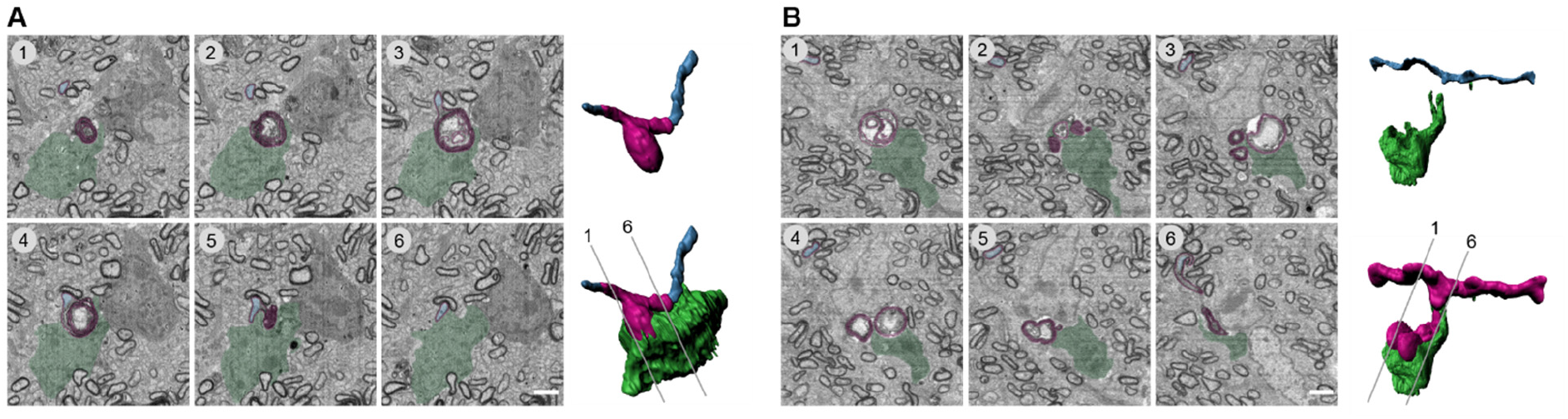
Microglia engulf and phagocytose developmental myelin pathology in the mouse CNS. (**A-B**) Serial block face scanning electron microscopy (SBF-SEM) of a P14 wild-type mouse optic nerve shows two examples of a microglia engulfing pathologic myelin (A and B). *Left:* Pseudocolored cross-sections (blue: axon, magenta: myelin, green: microglia). *Right:* 3D reconstruction. Numbering refers to cross-sections. Scale bars: 2 µm.

**Figure 2 – figure supplement 2.**
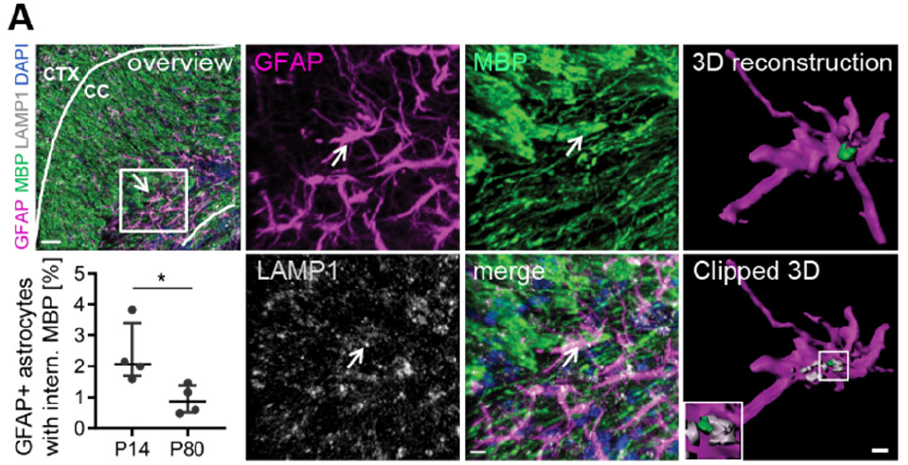
Astrocytes phagocytose myelin fragments in the mouse CNS. (**A**) Sections of P14 and P80 wild-type corpus callosum co-immunostained for MBP and LAMP1 together with GFAP for astrocytes. Clipped 3D view shows mbp-positive staining inside an astrocyte and in close association with lysosomes (arrows). Quantifications show the percentage of astrocytes with internalized MBP in P14 vs. P80 brains (n = 4 mice). Mann-Whitney test: *p<0.05. Data represent median ± interquartile range. Scale bars: 30 µm (overview), 7 µm (single channels), 3 µm (clipped 3D).

**Figure 3 – figure supplement 1.**
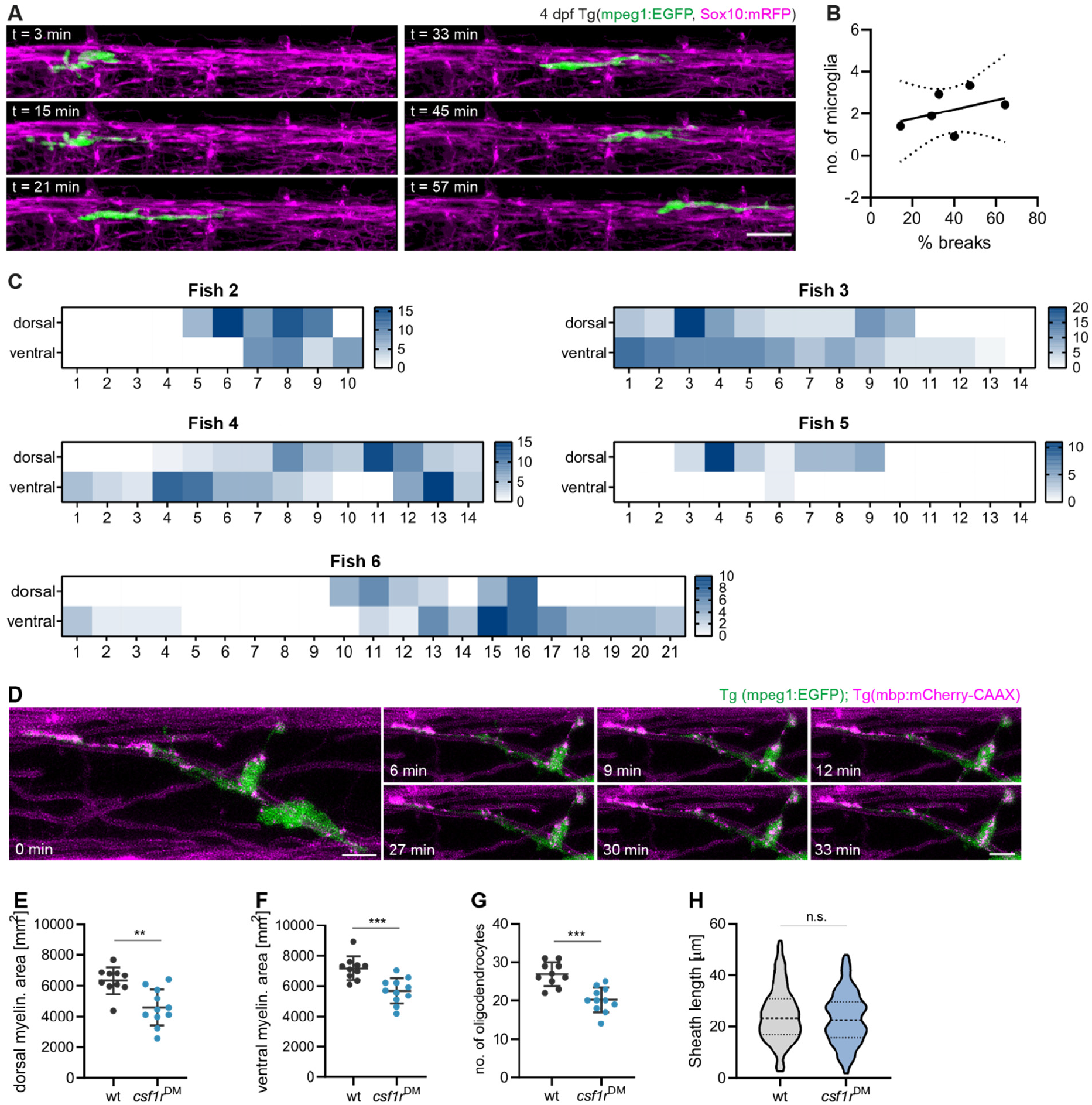
Screening behavior of microglia along myelin sheaths and analysis of myelination in csf1r^DM^ mutants. (**A**) Microglia motility along myelin sheaths was monitored in 3 min-acquisitions over 1 hour by confocal time-lapse imaging of a Tg(mpeg1:EGFP; sox10:mRFP) zebrafish spinal cord. (**B**) XY plot correlates the mean number of microglia present along the spinal cord with the mean percentage of frames in which microglia remained static. Data points represent individual larvae. Linear regression analysis: R^2^: 0.1669, p=0.4213. Dotted lines represent the standard error of the linear regression. (**C**) Heatmaps of microglial presence in quadrants of the dorsal and ventral spinal cord from 5 other fish (see Fig. 3F for Fish 1). (**D**) Confocal time-lapse imaging of motile mbp:mCherry-CAAX fragments within a microglia in a 7 dpf Tg(mpeg1:EGFP; mbp:mCherry-CAAX) spinal cord. Z-stacks were acquired every 3 min. Images show maximum projections. (**E-F**) Myelinated area in 10 dpf dorsal (E) and ventral (F) spinal cord (wt: n = 10, *csf1r*^DM^: n = 11 larvae). Student’s t test: p=0.0012 (E), p=0.0005 (F). (**G**) Number of oligodendrocytes in 10 dpf dorsal spinal cord (wt: n = 10, *csf1r*^DM^: n = 11 larvae). Student’s t test: p=0.0001. (**H**) Sheath lengths of 60 myelin sheaths each in the 10 dpf dorsal spinal cord of 3 larvae. Mann-Whitney test: p=0.3419. Data represent means ± s.d (E-G) and violin plots display median and interquartile range (H). Scale bars: 5 µm (D), 20 µm (A). See also Video 4.

